# LightAlign: a lightweight pairwise aligner for memory-constrained HiFi read assembly

**DOI:** 10.64898/2026.07.30.741934

**Authors:** Jian Liu, Jianwei Zhang

## Abstract

**Introduction:** Current *de novo* genome assembly tools often demand substantial memory resources, and their execution typically relies on high-performance computing (HPC) clusters. This dependency limits their use in resource-constrained settings. Furthermore, mainstream third-generation sequencing assembly and alignment tools usually require explicit detection of overlap regions between reads, a process that often entails significant computational and storage overhead.

**Results:** To address this issue, we developed LightAlign, a lightweight alignment tool for HiFi data that innovatively uses sequence-derived fuzzy features and reduces the peak memory usage during overlap detection.

**Conclusions:** When combined with miniasm, LightAlign generated bacterial draft assemblies while maintaining peak memory usage below 1 GB and completed overlap generation for the tested eukaryotic datasets within 1.88 GB RAM.

## 1 Background

*De novo* genome assembly is a fundamental step in genomic analysis, providing the sequence blueprint for downstream biological discovery. In OLC-based assembly, overlap detection is the starting point of the assembly process. Although exact alignment algorithms such as Needleman–Wunsch and Smith–Waterman provide rigorous approaches for sequence comparison, applying them directly to all possible read pairs is computationally prohibitive for large long-read datasets [1, 2]. Therefore, overlap detection is typically organized as a two-stage process: a filtration step first identifies candidate overlapping read pairs, and a more computationally expensive alignment step then verifies these candidate overlaps.

For third-generation sequencing reads, BLASR was among the first tools to reduce the cost of long-read alignment by using suffix-array or BWT-FM indexes to rapidly locate clusters of short exact matches as anchors, thereby restricting sparse dynamic programming and local refinement to candidate regions [3]. DALIGNER improved the efficiency of PacBio read overlap detection by partitioning reads into blocks, sorting extracted k-mers, and merging sorted k-mer lists to identify seed-sharing read pairs without exhaustively aligning all possible pairs [4]. MHAP reduced memory usage by compressing each read into a fixed-size MinHash sketch and using locality-sensitive hashing to estimate read similarity, so that only likely overlapping reads were subjected to coordinate estimation and further confirmation using shared sketch/minimizer evidence [5, 6]. Minimap and minimap2 further lowered the number of stored and compared seeds by selecting the minimum-hash k-mer from each sliding window, indexing these sparse minimizers, collecting exact minimizer hits between reads, chaining approximately collinear anchors, and extending only reliable chains when base-level alignments were required [7–9]. Strobemers improved seed informativeness by linking multiple non-adjacent sequence strobes into composite seeds, reducing sensitivity loss caused by sequencing errors or local sequence variation [10]. More recently, BLEND used SimHash-based fuzzy seed matching to recover approximate seed matches through hash lookup, reducing the need for strict exact seed matches while improving the sensitivity–speed trade-off in read overlapping and mapping [11].

Despite these advances, the memory and storage demands of long-read overlap detection can still exceed the capacity of consumer-grade hardware, forcing researchers to rely on large-scale computing clusters. This remains a major bottleneck that limits the broader use of genome assembly in small laboratories, start-ups, and educational institutions.

To address this challenge, we developed LightAlign, a memory-efficient pairwise aligner specifically designed for HiFi data. LightAlign introduces a novel strategy that uses compact fuzzy features instead of exact k-mer-derived seeds for candidate overlap screening, significantly reducing peak memory usage during sequence comparison. This design allows the tool to run efficiently on standard PCs and even portable laptops. When combined with the lightweight assembler miniasm [8], LightAlign enables high-quality draft genome assembly for a variety of species with minimal computational resources.

## 2 Materials and methods

### 2.1 LightAlign overview

The LightAlign workflow (Fig. 1) comprises five main steps. It begins with Input Standardization, where reads in FASTQ/FASTA format are imported and filtered by length. Next, Complementary Strand Generation computes the reverse complements for all retained reads. In Step 3, Fuzzy Feature Extraction derived fuzzy features from each read. Step 4, Correlated Read Pair Screening, is a key innovation of LightAlign—using fuzzy features rather than k-mer based methods to identify candidate overlapping read pairs. Finally, Step 5, Banded Dynamic Programming, verifies these candidate pairs and terminates alignments early when clear nucleotide-level mismatches are detected. After completing these five steps, LightAlign outputs mappings in pairwise mapping format (PAF), which are subsequently taken as input by miniasm for genome assembly.

**Fig 1.**
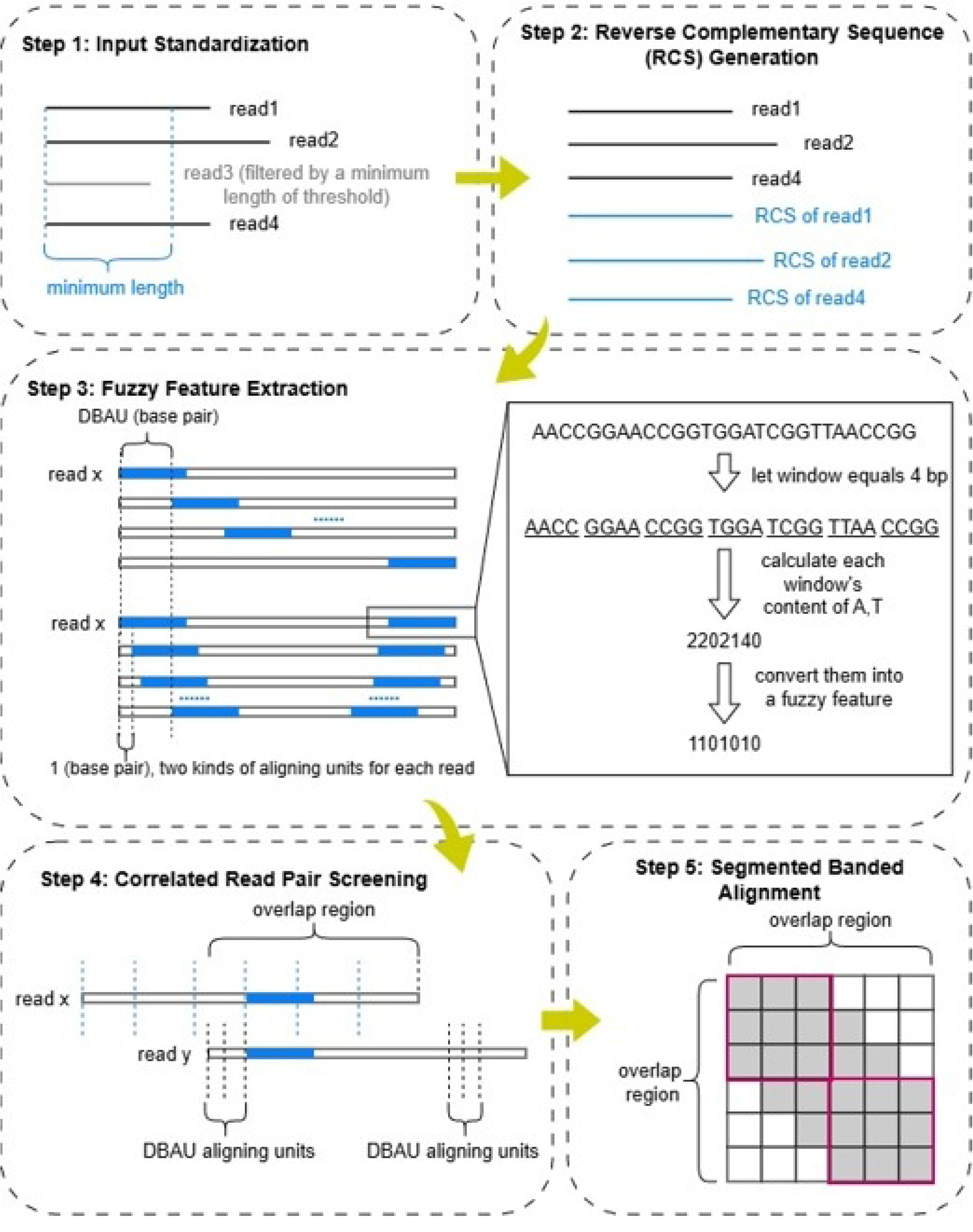
Overview of the LightAlign pipeline. Steps 1 and 2 perform data preprocessing, including read filtering based on minimum length, reverse complementary sequence generation, and input format standardization. Steps 3 and 4 handle candidate overlap detection—the core innovation of LightAlign. In Step 3, each blue rectangle represents an aligning unit. An aligning unit consists of multiple adjacent, non-overlapping windows. The fuzzy feature of an aligning unit is defined by fluctuation in AT content across these windows along the corresponding sequence segment. Specifically, if the AT content of a window is greater than or equal to that of the preceding window, it is encoded as 1; otherwise, it is encoded as 0. For each read (illustrated by *read x*), the two sets of aligning units are generated: one set is uniformly distributed along the read at fixed DBAU intervals, while the other is densely clustered at both the 5’ and 3’ ends, with the number of units at each end equals to the DBAU value. During pairwise alignment (Step 4), if two reads share an identical aligning unit (as illustrated by *read x* and *read y*), they are considered potentially overlapping and retained for further validation. Step 5 employs banded dynamic programming to confirm the overlaps identified in Step 4. The gray area represents the computed band, which is subdivided into equal-length segments to enhance cache efficiency. The gray portion within each red box indicates the computation performed per segment.

### 2.2 Data preprocessing

LightAlign begins by performing read-level preprocessing to ensure that only high-quality and sufficiently long sequence reads proceed to the overlap detection step (Step 1 in Fig. 1). Reads are first screened using a minimum length threshold, and any read shorter than this cutoff is discarded to prevent the generation of unreliable fuzzy features.

After length filtering, LightAlign computes the reverse complement of every retained read (Step 2 in Fig. 1). Both the original read and its reverse complement are stored on disk as separate entries so that overlap detection can be performed in a strand-agnostic manner. This design avoids the need of on-the-fly strand conversion during later alignment steps, thereby reducing runtime overhead. The preprocessing stage outputs a standardized dataset in FASTA format that serves as the input for fuzzy feature extraction.

### 2.3 Fuzzy-feature-based screening of read pairs

Read overlap detection typically begins by identifying potentially similar read pairs, followed by a more computationally intensive comparison using methods such as dynamic programming. Most current assemblers detect candidate overlaps by searching for shared seeds (usually short k-mers) between reads. These seeds are kept short to ensure that the vast majority of k-mers remain free from sequencing errors, as even a single base error can corrupt a k-mer and compromise overlap detection. If a considerable proportion of k-mers is affected by sequencing errors, the accuracy of this screening step is markedly reduced. In contrast, LightAlign identifies overlap candidates using fuzzy features, whose representation remains stable in the presence of a small number of sequencing errors, which allows the use of much longer sequence segments (over 1 kbp in our study) as seeds. Besides, LightAlign does not require collinearity checks across multiple seeds; each aligning unit typically represents several hundred or thousand base pairs in practice, thus the detection of a single identical aligning unit between two reads is sufficient for a read pair to pass screening.

Two key concepts are introduced in overlap detection process: the aligning unit and the distance between aligning units (DBAU). An aligning unit, LightAlign’s counterpart to the “seed” used in existing assemblers, refers to a series of fuzzy features derived from a specific segment of a nucleotide sequence. Its length is set to balance computational cost with detection sensitivity. The DBAU defines the spacing between the start positions of two aligning units. For each read, LightAlign generates two sets of aligning units. Although their window structures are identical, the two sets differ in distribution: one set is uniformly distributed along the read with a fixed DBAU interval, whereas the other is consecutively concentrated at both the 5′ and 3′ ends, with the number of units at each end equal to the DBAU value (Step 3 in Fig. 1). This design ensures that, when two reads share an overlap containing at least one aligning unit, a terminal aligning unit from one read—that is, an aligning unit from the second set located at either the 5′ or 3′ end—will always align with at least one corresponding unit from the other read, that is, an aligning unit from the first set. The DBAU is optimized to minimize the total number of aligning units per read, thereby reducing computational overhead. The detailed derivation of the aligning unit and DBAU parameters, along with the recommended values, is provided in Supplementary Note 2 in S1 File.

LightAlign creates fuzzy features by subdividing each aligning unit into windows and encoding the relative magnitudes of their calculated AT content (Step 3 in Fig. 1). Specifically, the AT content of each window is compared to that of the preceding window—except the first window is compared against zero. A window is encoded as 1 if its AT content is greater than or equal to the reference value; otherwise, it is encoded as 0 (Step 3 in Fig. 1). The significantly smaller informational size of the fuzzy features directly reduces LightAlign’s memory usage.

In candidate overlap screening (Step 4 in Fig. 1), LightAlign performs an all-vs-all comparison of fuzzy features based on the principle that overlapping fuzzy features may indicate true sequence overlap, whereas the absence of fuzzy feature overlap guarantees no sequence overlap.

In addition, LightAlign maintains relatively stable memory consumption across datasets of different sizes. Unlike minimap2, which keeps an index of the target sequences in memory, LightAlign uses a grouped-comparison strategy, loading only two read groups at a time for pairwise comparison. Thus, its peak memory usage is mainly determined by the group size rather than the total dataset size.

### 2.4 Banded dynamic programming alignment

The final step of the workflow is segmented banded alignment. LightAlign utilizes banded dynamic programming to align read pairs that have potential overlaps (Step 5 in Fig. 1). To improve cache efficiency and computational speed, the overlapping region between two reads is divided into multiple equal-length segments. Each segment is sequentially processed by banded dynamic programming, and the overall alignment score is obtained by summing the scores of all segments.

Before performing dynamic programming on a candidate read pair, the expected overlap length allows LightAlign to precompute the minimum score threshold required for a valid overlap, based on a predefined maximum error rate. Specifically, let *match*, *mismatch*, and *gap* denote the scores assigned to a match, mismatch, and a gap, respectively — where mismatch and gap are negative values, and the mismatch penalty is greater in magnitude than that of the gap. Given the observed overlap length (*actualOverlap*) and the maximum allowed error rate (*errorRate*), the score threshold is calculated as:

*threshold* = (1 - *errorRate*) × *match* × *actualOverlap* + *errorRate* × *mismatch* × *actualOverlap*.

During the segmental alignment, LightAlign continuously tracks the current cumulative score (*currentScore*) and the total length of processed regions (*processedLength*). After completing each segment, the program evaluates the following early-termination condition:

*currentScore* + (*actualOverlap* - *processedLength*) × *match* < *threshold*.

If this condition is met, the alignment for that read pair is immediately terminated to avoid unnecessary computation. Despite these optimizations, Step 5 remains the most computationally demanding and time-consuming stage of the entire LightAlign workflow.

### 2.5 Benchmarking LightAlign for HiFi read overlap and assembly

We benchmarked LightAlign using simulated reads and seven public HiFi datasets, including four prokaryotic and three eukaryotic datasets. For prokaryotic datasets, we compared the LightAlign–miniasm pipeline with the minimap2–miniasm pipeline using identical miniasm commands. All LightAlign runs were performed on a personal computer equipped with a 6-core, 12-thread CPU. Miniasm v0.3 [8] and minimap2 v2.28 [12] were used in all benchmarking experiments. Minimap2 was run with 12 threads, and the -c option was enabled to generate CIGAR strings for consistent comparison. This option increases the computational workload of minimap2 but is not expected to materially affect its peak memory usage.

Because minimap2 does not include a preset specifically designed for overlap detection of PacBio HiFi reads, two minimap2 parameter settings were evaluated. The first setting used the default minimap2 preset for PacBio all-vs-all long-read alignment, -x ava-pb. The second setting was adjusted according to the high base-level accuracy and low sequencing error rate of HiFi reads. Specifically, the k-mer size, window size, minimum chaining score, maximum gap, bandwidth, and minimum number of secondary chains were modified using the following parameters: -x ava-pb -k19 -w10 -m100 -g2000 -r500 -n3. Detailed parameter settings and command lines are provided in Supplementary Notes 2 and 3 in S1 File.

## 3 Results

### 3.1 Evaluation of LightAlign’s overlap-detection sensitivity

First, 30× simulated reads were generated with Badread [13] from the *Saccharomyces cerevisiae* R64 reference genome GCF_000146045.2 with the command provided in Supplementary Note 1 in S1 File. Ground-truth overlaps between reads were then defined from their known genomic coordinates, using a minimum overlap length of 750 bp, consistent with the aligning-unit length parameter used by LightAlign in this test (-l 750 bp). Then we compared the overlap-detection performance of LightAlign and minimap2 on the simulated reads and used the overlaps produced by each tool as input for miniasm assembly. Finally, contigs generated by the LightAlign–miniasm pipeline were aligned to the reference genome GCF_000146045.2 to assess assembly accuracy.

We compared LightAlign with the two minimap2 settings described in Methods and used the overlaps produced by each tool as input for miniasm assembly. Overall, LightAlign and minimap2 exhibited distinct overlap detection characteristics on the simulated reads. Minimap2 achieved a higher recall; however, its lower precision indicated that the detected overlap relationships contained a larger number of false positives. In contrast, although LightAlign showed a relatively lower recall, it achieved higher precision and better overall F1 performance, where the F1 = 2 × precision × recall / (precision + recall) (Table 1). Although the LightAlign-miniasm pipeline yielded a slightly lower N50 than the minimap2-miniasm pipeline, it produced fewer contigs, closer to the 17 chromosomes of the reference genome, and did not generate the short contigs observed in the minimap2-based assemblies (Table 2).

**Table 1.** Overlap-detection performance of LightAlign and minimap2 on simulated reads.

| Tools | Parameters | Recall | Precision | F1 | Orientation accuracy |
| --- | --- | --- | --- | --- | --- |
| LightAlign | l = 750 bp | 0.78 | 0.96 | 0.86 | 1.00 |
| minimap2<br>(default) | default | 1.00 | 0.48 | 0.65 | 1.00 |
| minimap2<br>(adjusted) | adjusted | 1.00 | 0.49 | 0.65 | 1.00 |

**Table 2.** Assembly performance of LightAlign–miniasm and minimap2–miniasm pipelines on simulated reads.

| Tools | Contigs | Total length (Mb) | Max length (Mb) | N50 (Mb) |
| --- | --- | --- | --- | --- |
| LightAlign | 18 | 11.26 | 1.49 | 0.78 |
| minimap2<br>(default) | 20 | 12.11 | 1.53 | 0.92 |
| minimap2<br>(adjusted) | 20 | 12.11 | 1.53 | 0.92 |

The resulting contigs showed clear collinearity with the reference genome, supporting the reliability of the LightAlign-based assembly, and no obvious secondary alignment signals were observed in the dot plot (Fig 2). These results suggest that, although LightAlign does not match existing tools in terms of recall, it can still support the generation of relatively high-quality assembly results.

**Fig 2.**
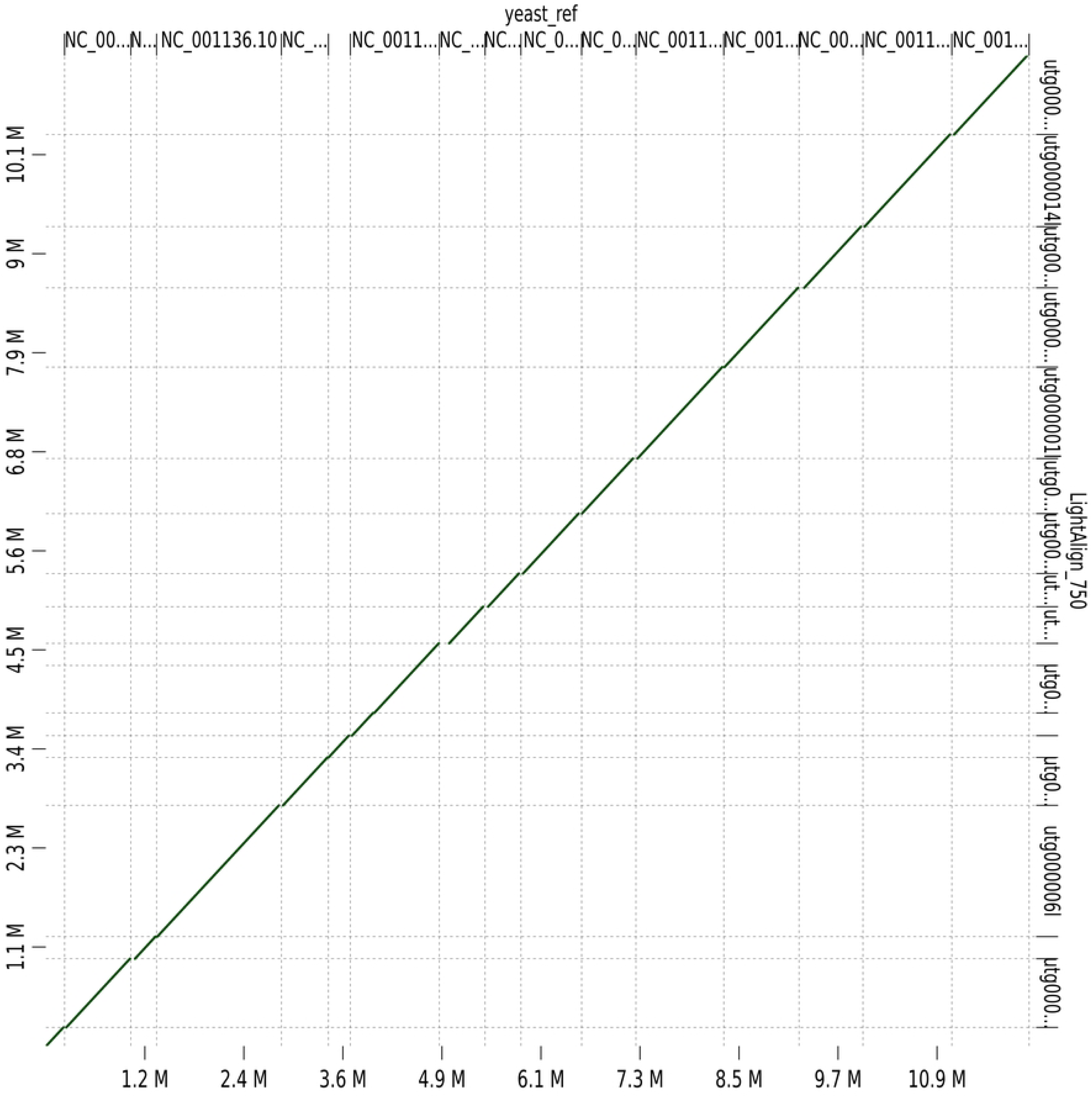
Dot plot comparison between contigs assembled from 30× simulated reads generated based on the reference genome GCF_000146045.2 using the LightAlign-miniasm pipeline and the corresponding reference genome GCF_000146045.2. The clear collinear alignment pattern and the absence of substantial secondary alignments indicate high assembly quality.

The lower recall of LightAlign is mainly attributable to the nature of its fuzzy feature, which is defined by the relative changes in AT content between adjacent windows. Insertion and deletion (indel) errors can shift window boundaries and thereby alter the AT-content fluctuation pattern across multiple downstream windows, reducing the consistency of the fuzzy feature. Overlaps with relatively frequent indel errors may therefore be missed. By contrast, substitution errors affect only the AT content within individual windows and are less likely to change the relative AT-content relationships between adjacent windows, making LightAlign more robust to this type of error. In addition, the minimum overlap detectable by LightAlign is defined by the length of a single aligning unit -- typically several hundred to over one thousand base pairs -- whereas minimap2 uses a default minimum overlap threshold of 100 bp. This inherent difference in parameter settings also naturally results in fewer overlaps being identified by LightAlign.

Despite lower overlap-detection sensitivity than minimap2, LightAlign achieved comparable assembly performance on real datasets, as reflected by N50 and total assembly length. One possible explanation is that not every detected overlap contributes directly to the final assembly. Miniasm constructs a string graph from suffix–prefix overlaps and simplifies the graph by removing transitive and other redundant overlap relationships [8]. Consequently, some overlaps missed by LightAlign may have been redundant for the assembly paths recovered in the tested datasets. This may explain why the lower overlap-detection recall of LightAlign did not necessarily result in markedly poorer draft-assembly statistics. However, this observation is dataset-dependent and does not imply that reduced overlap recall is generally inconsequential, particularly for repetitive, heterozygous, or structurally complex genomes.

### 3.2 Performance of LightAlign on prokaryotic datasets

We evaluated LightAlign on seven public HiFi datasets, including four bacterial and three eukaryotic datasets (Table 3). For the bacterial datasets, we compared the LightAlign– miniasm pipeline with minimap2–miniasm pipelines generated using the two minimap2 parameter settings described in Methods. In all cases, miniasm was run with identical commands on the PAF outputs produced by each aligner. Detailed parameter settings and command lines are provided in Supplementary Notes 2 and 3 in S1 File.

**Table 3.**
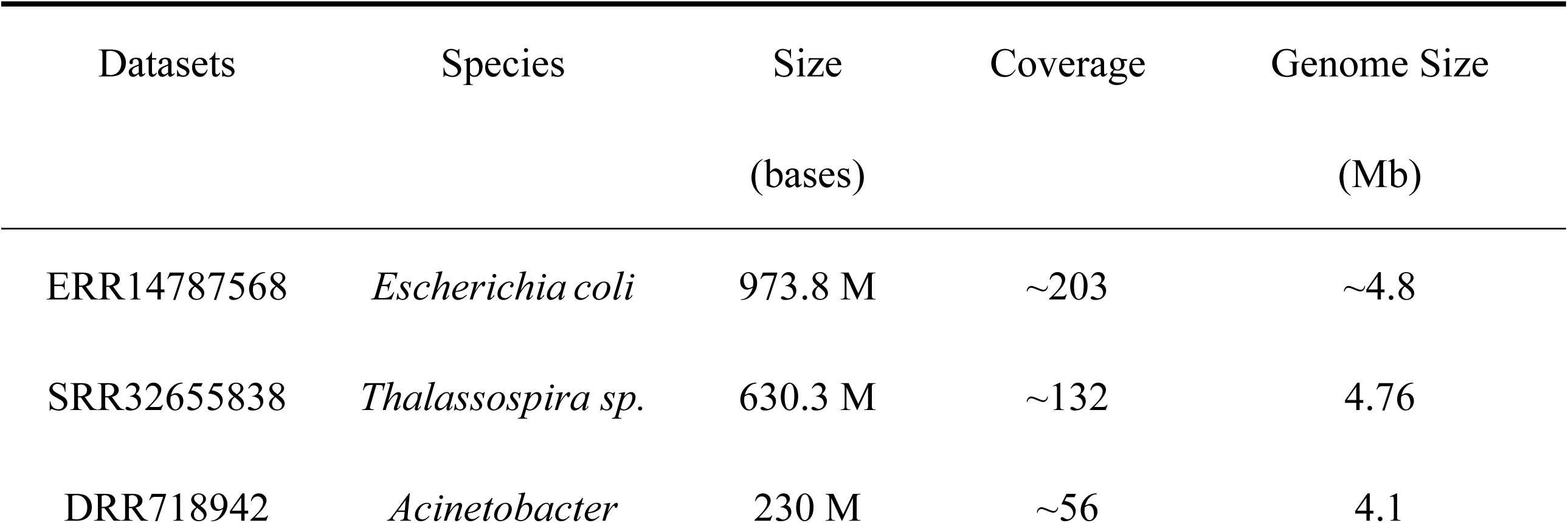
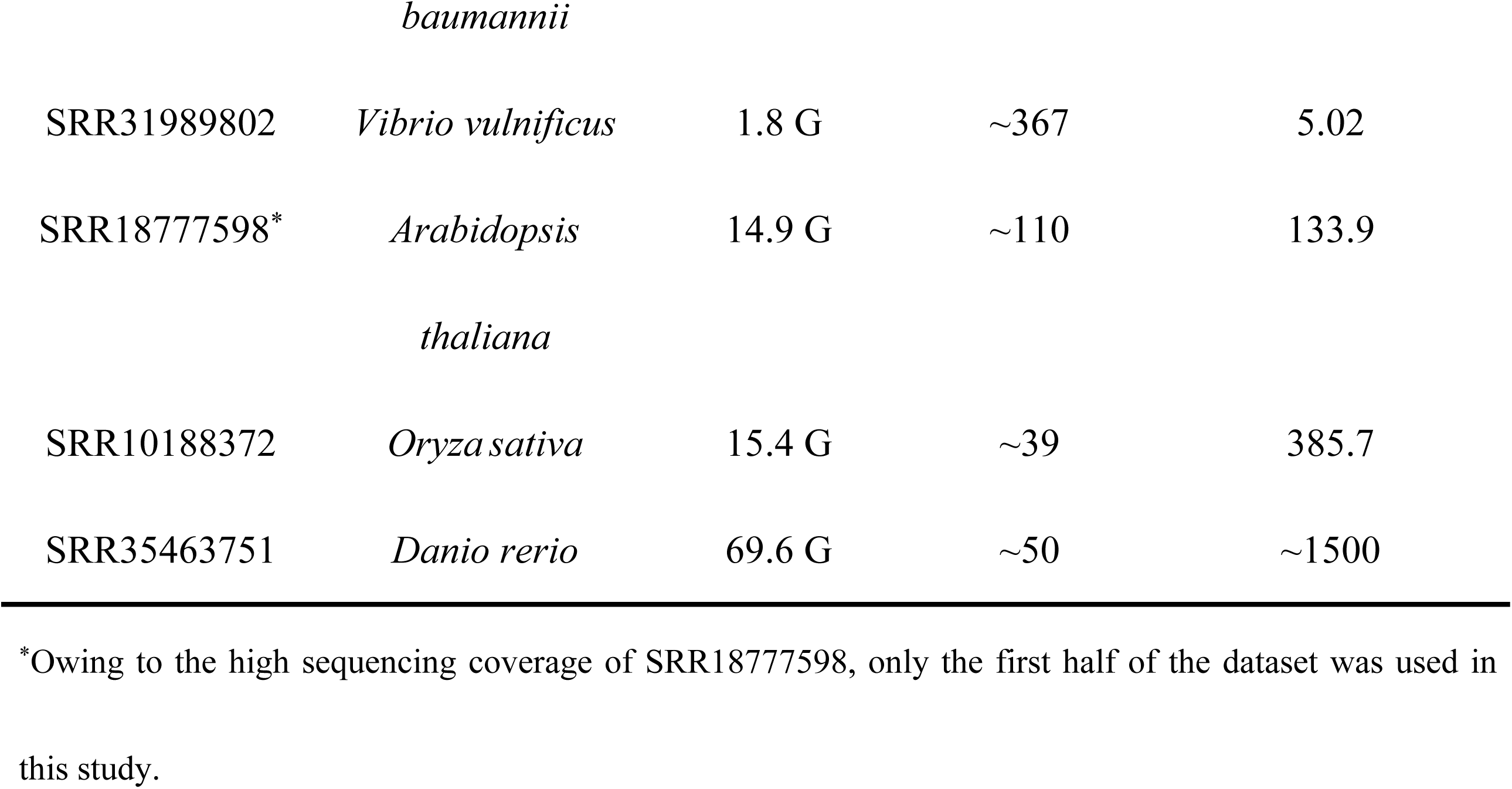
Datasets used for aligner performance evaluation.

Benchmarking results show that LightAlign was 1.5x–4.3x slower (on average 2.7x) than minimap2, but its overall runtime remained within practical limits for bacterial genome assembly. In contrast, LightAlign showed a substantial memory advantage, using 3.1x–71.9x less memory (on average 26.3x) than minimap2. This reduction in memory usage became more pronounced on larger-scale datasets. When paired with miniasm, the LightAlign– miniasm pipeline produced assemblies with N50 values comparable to, and in some cases higher than, those generated by the minimap2–miniasm pipeline across all four datasets (Fig 3, Supplementary Table 1 in S1 File). In addition, contigs generated by LightAlign and minimap2 showed good collinearity (Supplementary Figs 1–4 in S1 File).

**Fig 3.**
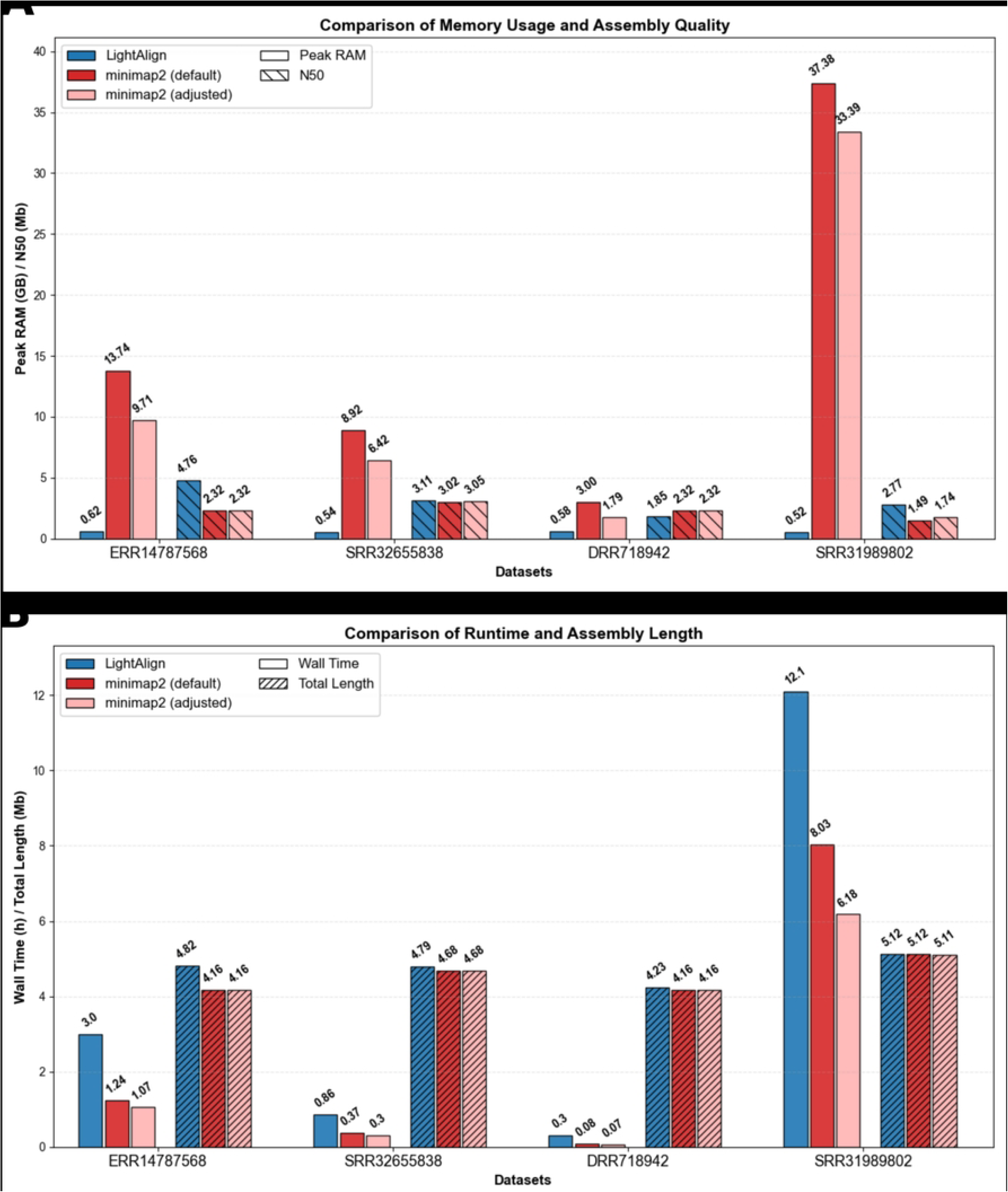
Computational resource usage and assembly performance of LightAlign compared with minimap2. (A) Peak RAM and contig N50 values for LightAlign and minimap2 across four bacterial datasets. (B) Runtime and assembly length for LightAlign and minimap2 across four bacterial datasets.

The memory saving arises from two design choices in the overlap-generation stage. First, during the pairwise comparison stage, the fuzzy features used by LightAlign require only a small amount of data to represent reads. For example, when window = 30 bp, DBAU = 87, length of an aligning unit = 900 bp, a 15 kb read converted into fuzzy features occupies only approximately 17% of the memory required to store the original nucleotide sequence within the program. Second, it uses a grouped-comparison strategy in which only two read groups are loaded into memory at a time. Therefore, its peak memory usage is governed mainly by the group size and feature representation, rather than by the total number of reads in the dataset. This explains why LightAlign maintains low and relatively stable memory usage across datasets of different scales.

### 3.3 Performance of LightAlign on eukaryotic datasets

To evaluate the performance of LightAlign on eukaryotic genomes, we assembled three HiFi sequencing datasets from *Arabidopsis thaliana*, *Oryza sativa*. and *Danio rerio* (Table 3) with LightAlign-miniasm pipeline. However, minimap2 did not produce results for the eukaryotic datasets because its memory usage exceeded the computational resource limit set in this study (60 GB).

The memory usage of LightAlign remained low at 0.74 GB, 0.81 GB, and 1.88 GB across three datasets. The resulting contigs were then aligned to their respective reference genomes Col-PEK, MH63RS3 and GRCz12tu using D-GENIES [14]. While the dot plots confirm high similarity to the references (Supplementary Figs. 5-7 in S1 File), the assemblies show reduced contiguity compared to bacterial ones, with N50 lengths of 0.91 Mb, 0.63 Mb, and 0.43 Mb (Supplementary Table 2 in S1 File). This limitation is likely attributable to the fact that miniasm is not specifically designed for assembling highly complex eukaryotic genomes [8].

## 4 Discussion

Across all bacterial datasets, LightAlign consistently maintained peak memory usage below 1 GB when processing HiFi sequencing data. While users with substantial computational resources may achieve the shortest runtimes for bacterial genome assembly by combining minimap2 with assemblers such as miniasm, LightAlign provides a practical and reliable alternative for memory-constrained or resource-limited environments. For the eukaryotic datasets, the LightAlign-miniasm pipeline generated reasonably accurate yet less contiguous contigs compared to bacterial results. This limitation likely stems from miniasm not being designed for assembling highly complex eukaryotic genomes. Notably, LightAlign maintained extreme memory efficiency on eukaryotic datasets, and opens the door to *de novo* eukaryotic genome assembly on consumer-grade hardware.

Sequencing errors like insertions and deletions (indels) may shift window boundaries and disrupt fuzzy feature correspondence, constraining the current applicability of LightAlign primarily to high-fidelity (HiFi) sequencing data, which exhibit very low indel rates. Building upon LightAlign’s capability for ultra-low memory usage on both prokaryotic and eukaryotic datasets, future work will focus on developing an assembler that similarly prioritizes memory efficiency while offering enhanced capability to handle highly repetitive genomes. In addition, to further lower hardware barriers at the data acquisition stage, a dedicated low-memory error-correction tool should be developed for Oxford Nanopore Technologies (ONT) data, which are substantially more cost-effective than HiFi sequencing. The corrected reads generated by this tool would serve as input for LightAlign, thereby extending its applicability to a broader range of sequencing platforms.

## 5 Conclusions

LightAlign introduces a fuzzy-feature-based overlap-generation strategy that substantially reduces peak memory usage for HiFi read assembly. Combined with miniasm, it enables bacterial genome assembly under 1 GB RAM and supports eukaryotic genome assembly on consumer-grade hardware. These results demonstrate that memory-efficient overlap generation can lower the computational barrier for long-read genome assembly in resource-limited settings.

## 6 Acknowledgements

We thank the National Key Laboratory of Crop Genetic Improvement and the Supercomputing Center at Huazhong Agricultural University (HZAU) for providing the bioinformatics computing platform in this study.

## Supporting information

**S1 File. Supplementary methods, parameter settings, benchmark results, and dot plots for LightAlign.** This file contains Supplementary Notes 1–3, Supplementary Tables 1–2, and Supplementary Figs 1–7.

